# Genetically modified IL-2 bone marrow-derived myeloid cells reprogram the glioma immunosuppressive tumor microenvironment

**DOI:** 10.1101/2022.10.19.511786

**Authors:** Alessandro Canella, Matthew Nazzaro, Sakthi Rajendran, Claire Schmitt, Abigail Haffey, Giovanni Nigita, Diana Thomas, Haley Wrightnour, Paolo Fadda, Elaine R. Mardis, Timothy P. Cripe, Prajwal Rajappa

**Affiliations:** The Steve and Cindy Rasmussen Institute for Genomic Medicine, Nationwide Children’s Hospital, Columbus, OH, USA; Department of Cancer Biology and Genetics, Comprehensive Cancer Center, The Ohio State University, Columbus, OH, USA; Department of Pathology and Laboratory Medicine, Nationwide Children’s Hospital, Columbus, OH, USA; Animal Resource Core, Nationwide Children’s Hospital, Columbus, OH, USA; Genomics Shared Resource, The Ohio State University and James Cancer Hospital, Columbus, OH; Center for Childhood Cancer, The Abigail Wexner Research Institute, Nationwide Children’s Hospital, Columbus, OH, USA

## Abstract

Gliomas are the most prevalent type of brain tumors and one of the leading causes of cancer-related death in the adolescent and young adult population (AYA). Two-thirds of glioma AYA patients are affected by low-grade gliomas (LGGs), but there are no specific treatments. Therefore, a percentage of LGG patients experience tumor relapse and malignant progression to high-grade glioma which leads to fatal outcomes. In part, malignant progression is potentiated by the immunosuppressive stromal component of the tumor microenvironment (TME) underscored by M2-macrophages and a paucity of cytotoxic T cells. As a result, first-line immunotherapies have failed to improve outcomes for patients with progressive high-grade gliomas. Here, we report the efficacy of an in vivo approach that demonstrates the potential for a novel cell-mediated innate immunotherapy designed to abrogate immunosuppressive mechanisms within the glioma TME and enhance the recruitment of activated effector T cells. A single dose of engineered bone marrow-derived myeloid cells that release Interleukin-2 (GEMys-IL2) was used systemically to treat mice with LGG tumors systemically. Our results demonstrate that GEMys-IL2 efficiently crossed the blood brain barrier (BBB), infiltrated the glioma microenvironment, and reprogrammed the infiltrating immune cell composition and transcriptome. In addition, GEMys-IL2 impaired tumor progression and extended survival in a LGG immunocompetent mouse model. In conclusion, we demonstrated that GEMys-IL2 have a therapeutic effect in vivo, thus supporting its potential application as a novel immunotherapy that warrants further investigation.

## INTRODUCTION

In the United States, approximately 60,000 new cases of brain tumors have been diagnosed from 2012 to 2016, and nearly 26% were gliomas [1, 2]. Tumors of the central nervous system (CNS) are one of the most prevalent solid tumors and the third leading cause of cancer-related death in adolescents and young adult (AYA) patients (from 15 to 39 years of age)[1, 3]. More importantly, there is no established therapeutic protocol for treating gliomas in AYA patients. At the time of diagnosis, two-thirds of AYA patients affected by glioma are histologically low-grade (LGG, WHO grade I and II), and one-third are high-grade gliomas (HGG, WHO grade III and IV) [4, 5]. LGG represents a heterogeneous group of primary tumors derived from neuroglial cells [6], and in LGG AYA patients the most frequent histological types are astrocytoma (pilocytic grade I, diffuse grade II) and oligodendroglioma (grade II) [7]. In these patients, the five-year progression-free survival (PFS) after total resection is above 90%, but it significantly decreases to 55% in those with residual disease [8]. Despite the favorable response of the patients to the standard of care, which consists of total resection followed by chemo-radiotherapy, a significant number of AYA patients experience tumor relapse, associated with a faster progression to HGG and a significant reduction of survival to 1-3 years post-relapse [9, 10].

In the last decade, efforts have started to characterize the tumor biology and genetic heterogeneity of gliomas, but the molecular processes by which tumors relapse, and transform into high-grade disease, remain for the most part, unknown. The glioma microenvironment is a pro-tumoral tridimensional architecture mainly constituted by a network of cancer cells, stromal cells (immune cells, fibroblasts, endothelial cells), blood vessels, and extracellular matrix. The stromal components of the TME support homing and proliferation of the cancer cells through the release of pro- or anti-inflammatory chemokines, exosomes, pro-angiogenetic factors, pro-survival chemokines, and growth factors [11]. Moreover, the TME assists in the development of immune escape mechanisms that protect the cancer cells during tumor progression by hindering cancer immune surveillance, led by innate and adaptive immune cells. Cancer cells facilitate the immunosuppressive reprogramming of myeloid cells infiltrating the TME [12] through the release of cytokines (IL4, IL10, and TGFβ), restricting the recruitment and activation of cytotoxic effector immune cells, such as cytotoxic T lymphocytes (CD3+CD8+, CTLs), dendritic (DCs) and natural killer (NK) cells [13, 14]. Interestingly, glioma tumor progression is characterized by a significantly increased infiltration of myeloid cells and a reduction of infiltrating cytotoxic T and NK cells in the TME [15, 16]. In fact, approximately 50% of the immune cells in the TME are myeloid cells, either resident (microglia) or recruited from the bone marrow (bone marrow-derived myeloid cells - BDMCs) [16, 17]. Myeloid cells in the TME can have a range of pro-activating (M1-like) or immunosuppressive (M2-like) phenotypes [18], and the balance between M1-like and M2-like myeloid cells, together with the infiltration of immunosuppressive regulatory T cells (Tregs) in the tumor niche, are clinically prognostic [19, 20]. To that end, increased tumor infiltration of Tregs and M2-like macrophages have been associated with reduced sensitivity to drug treatments, including immunotherapies, and with a significant reduction of PFS in glioma patients [21]. In the last decade, immunotherapy was considered a major advance for the treatment of several tumor types, but it has shown very limited efficacy in glioma [22]. This failure is thought to be mainly due to the presence of the blood brain barrier (BBB), which hampers the ability of these drugs to reach the TME, and to immune-escape mechanisms mediated by cancer cells and immunosuppressive myeloid cells [23, 24].

The reduction of chemo attractant and immune stimulatory molecules within the TME, and the immunosuppression promoted by myeloid and Treg cells, appear to hamper the recruitment and activation of intratumoral cytotoxic immune cells such as CTLs and NK cells [25, 26]. Interleukin-2 (IL2) is a 15.5KDa cytokine secreted by activated T lymphocytes and one of the master activators of naïve T cells [27]. The interaction between IL2 and its receptor triggers JAK-STAT and mTOR signaling pathways associated with cellular growth, proliferation, and cell cycle progression to drive the cellular differentiation from the naïve to the effector T cell phenotype. On the other hand, excessive or prolonged IL2 stimulation of T lymphocytes can affect the activation status and lead to T cell exhaustion or inactivation [28].

In this study, we designed and pre-clinically evaluated a novel therapeutic concept to deliver bone marrow-derived myeloid cells (GEMys), genetically engineered for the stable expression and release of interleukin-2 (IL2) within the TME, as a means to promote the recruitment and activation of CTLs against tumor cells. We demonstrated that the intravenous injection of a single dose of GEMys-IL2 in LGG mice was sufficient to rapidly recruit and activate more CTLs against tumor cells in the TME. The GEMys released IL2 in the glioma microenvironment, thus promoting the trafficking and activation of cytotoxic T cells and altering the composition and transcriptome of the immune cells. Importantly, the GEMys-IL2 treatment in vivo delayed tumor progression and significantly improved the overall survival in LGG bearing animals (OS).

## METHODS

### Additional methods are described in the Supplementary file

#### Generation of genetically engineered myeloid cells for the stable expression of IL2

The generation in vitro of control GEMys (empty vector, EV), or GEMys for the stable expression and release of IL2, was performed as previously described [29]. Briefly, bone marrow cells were isolated from mice not bearing tumors (NTV-a;Ink4a^-^Arf^+/-^;PTEN^fl/fl^;LSL-Luc), and the hematopoietic progenitors were isolated by magnetic negative selection (Mouse Hematopoietic Progenitor Cell Isolation Kit, StemCell Technologies), and cultivated in SFEM II media supplemented with 50ng/ml of murine IL6, SCF, and FLT3-L (StemCell Technologies) and 1% penicillin/streptomycin to drive the differentiation to mature myeloid cells. After 2 hours, cells were infected with lentiviral particles (MOI=100) in the presence of Polybrene 10μg/ml (Sigma-Aldrich). The lentiviral particles were generated by co transfection in Lenti-X™ 293T cells (Takarabio) of lentiviral packaging kit (Origene) and plasmids for the expression of IL2-GFP+, or the empty vector-GFP+, according to the manufacturer’s protocol (Origene). The lentivirus concentration was quantified by Lenti ELISA Kit (Origene). After 16h, the cells were scraped, washed, and seeded in fresh media. Three days post-infection, the cells were tested for the expression of IL2 by RT-qPCR, and for the release of IL2 by ELISA (R&D Systems). Infected myeloid cells were selected by the addition of puromycin 1μg/ml for 2 weeks, and the GFP and IL2 expression were monitored weekly by flow cytometry (LSR Fortessa, BD Biosciences), fluorescent microscopy (EVOS, AMG), RT-qPCR (Applied Biosystems), and ELISA (R&D Systems).

#### Mouse model

NTV-a;Ink4a^-^Arf^+/-^;PTEN^fl/fl^;LSL-Luc and the RCAS system were adopted to undergo gliomagenesis in vivo [30]. DF-1 chicken fibroblasts transfected with RCAS-PDGF and DF-1 cells transfected with RCAS-Cre were maintained in DMEM with 10% fetal bovine serum (Gibco). Heterozygous NTV-a;Ink4a^-^Arf^+/-^;Pten^fl/fl^;LSL-Luc were generated by crossing NTV-a (WT) mice with NTV-a;Ink4a^-^Arf^-/-^;Pten^fl/fl^;LSL-Luc mice. 0-2 days after birth, heterozygous pups were injected with 1 μL DPBS (Gibco) containing the two fibroblast cell lines (10^5^ cells/kind). Injections were executed in the right hemisphere of the brain parenchyma [31]. When mice were 4 weeks old, D-luciferin (Perkin Elmer) was administered by IP injection, and tumor burden was monitored by bioluminescence imaging (IVIS Spectrum, Perkin Elmer).

#### Evaluation of the treatment in the TME

Gliomagenesis was induced by injection of DF1-CRE and DF1-PDGF cells in NTV-a;Ink4a^-^Arf^+/-^;PTEN^fl/fl^;LSL-Luc pups at 0 to 2 days of age, as already described. At 4 weeks old, tumor burden was evaluated by IVIS, followed by mice randomization and separation into 3 treatment groups. At day 28, mice were retro-orbitally injected with 80μl of DPBS (vehicle), or with 8 million of GEMys-EV or GEMys-IL2. Mice were monitored twice a day and euthanized at day 5 or day 7 post treatment, followed by the isolation of the brain and tissue processing for further investigation.

#### Survival studies

Gliomagenesis was induced by injection of DF1-CRE and DF1-PDGF cells in NTV-a;Ink4a^-^Arf^+/-^;PTEN^fl/fl^;LSL-Luc pups at 0 to 2 days of age, as previously described. At 4 weeks of age, tumor burden was evaluated by IVIS, followed by mice randomization and separation in 3 treatment groups. At day 28, mice were retro-orbitally injected with 80μl of DPBS (vehicle), 8 million of GEMys-EV or of GEMys-IL2. Mice were monitored twice a day and euthanized at the beginning of decline, with symptoms such as lethargy, weight loss, macrocephaly, seizure, hyperactivity, or abnormal behavior.

#### Tissue harvesting

**Bone Marrow**: Femurs and tibias were dissected in sterility from NTV-a;Ink4a^-^ Arf^+/-^;PTEN^fl/fl^;LSL-Luc mice not bearing tumors 4 weeks of age. Bone marrow was extracted by centrifugation at 10,000 x g for 30 seconds. **Spleen:** Spleens were dissected from NTV-a;Ink4a^-^ Arf^+/-^;PTEN^fl/fl^;LSL-Luc mice without tumors at 4 weeks of age. **Peripheral Blood:** At endpoint, mice were anesthetized with an isoflurane vaporizer (Kent Scientific), and cardiac puncture was performed to collect 400μl of peripheral blood. Blood was transferred into microtainer EDTA blood collection tubes (BD), and animals were immediately euthanized. Blood samples were centrifuged at 2,000 x g for 10 minutes at +4°C, serum was collected and immediately analyzed by ELISA, or stored at −80°C. **Brain:** Whole brains were dissected from mice at endpoint and fixed in 10% buffered formalin phosphate for 24 hours for histology, or tumor area was extracted from the whole right hemisphere with a scalpel, dissociated to isolate immune cells of the glioma microenvironment for further investigation of the cellular composition by flow cytometry, or analysis of the transcriptome by RT-qPCR, and by Nanostring.

All animal studies were approved by Nationwide Children’s Hospital Institutional Animal Care and Use Committee (protocol AR19-00146).

#### Statistical analysis

All the results were generated by multiple independent observations and expressed as the mean ± standard deviation (SD). Overall survival was defined as the time from the injection of cells to generate gliomagenesis to the time of the endpoint and measured by Kaplan-Meier curve. Survival analyses were performed using Log-rank (Mantel-Cox) test. All reported p-values were calculated by unpaired two-tailed t-test, and analyses were performed using GraphPad (Prism). Experiments generated by at least three independent experiments with a p-value<0.05 were considered significant.

## RESULTS

### Glioma malignant progression effects the recruitment of T cells in the TME

Recent studies have demonstrated that glioma cells reprogram immune cells that are infiltrating the tumor microenvironment to trigger drug resistance mechanisms [32, 33] and develop an immunosuppressive phenotype, blocking the recruitment and activation of cytotoxic T and NK cells [25]. In our study, Ingenuity Pathway Analysis (IPA) of single cell RNA sequencing data (scRNAseq) obtained from the tumor infiltrating lymphocytes (TILs) in HGGs compared with LGGs, was used to predict upstream regulators, altered canonical pathways, and disease-related biological functions. As shown in Figure 1A-C and Supplementary Figure 1B-C, the analysis highlighted the effect of tumor progression on the immunosuppression of the T cell compartment. Indeed, we found a strong downregulation of IFNγ and a reduction of T cells trafficking (Fig. 1A). Moreover, the gene enrichment analysis by disease and function showed a strong decrease of genes involved in activation, migration, and trafficking of T lymphocytes (Fig. 1B, red arrows and Suppl. Table 1), resulting in decreased tumor infiltration by T lymphocytes, and in particular by CD8+ T cells (Fig. 1C, red arrows, and Suppl. Table 2). In addition, T cell receptor signaling was significantly downregulated (Suppl. Fig. 1B, z-score= −2.236, p=0.0141). This pathway is critically important for effector T cell proliferation and activation, and it is crucial for naïve T cell differentiation into effector T cells [34]. Furthermore, the pathway responsible for apoptosis of the target cells by cytotoxic T lymphocytes was significantly dysregulated during the transition from low- to high-grade glioma (Suppl. Fig. 1C, p=0.000126).

**Figure 1.**
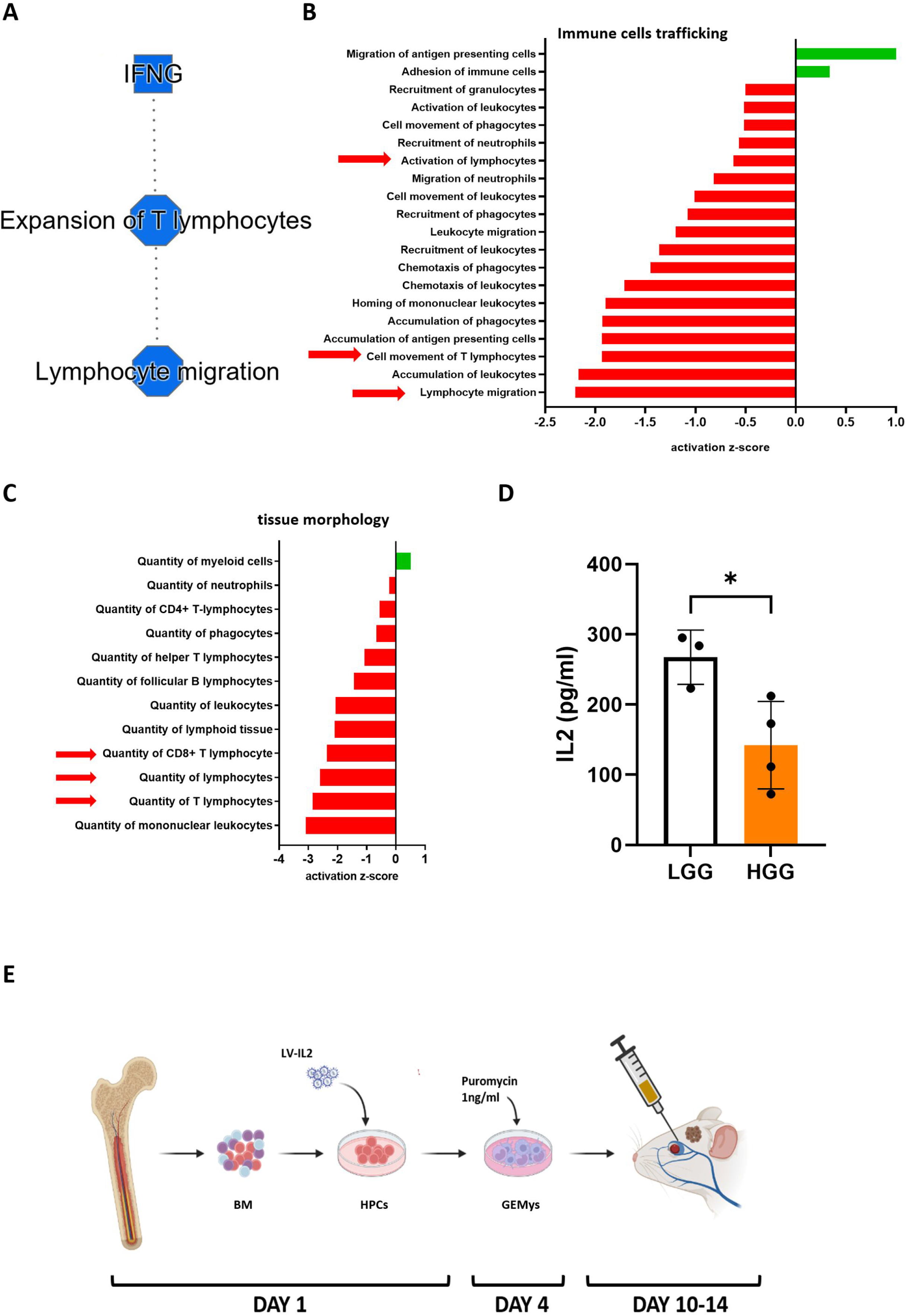
The Glioma progression from the low- to the high-grade affect cytotoxic T cells trafficking and activation in vivo. **A)-C)** IPA analysis of the differential gene expression of TILs isolated from RCAS/t-va LGG and HGG. The data were generated by scRNAseq (n=3). **A)** IPA graphical summary of the most important downregulated pathways **B)** IPA analysis of disease and biological function of the TILs trafficking (p<0.05). **C)** IPA analysis of disease and biological function of the TILs tissue morphology (p<0.05). **D)** Analysis by cytokine array of the intratumoral protein expression of IL2 in RCAS/t-va LGG, and HGG animals (n=3-4).). Unpaired two-tailed student’s t test calculated statistical significance. *P<0.05. **E)** Summary of the design of the novel cell-based immunotherapy (biorender.com).

In line with our scRNAseq findings, which revealed a significant reduction of T cell activation in the TME, the cytokine analysis of the glioma microenvironment during malignant progression from low- to high-grade glioma showed a significant downregulation of IL2, CCL21, CX3CL1 in the tumor milieu, which are associated with T cell recruitment, proliferation and activation (Fig. 1D, Suppl. Fig. 1D,E) [27, 35]. Based on these results, we evaluated the ability of engineered bone marrow-derived myeloid cells to reprogram the immunosuppressive myeloid cell rich TME, to recruit anti-cancer immune cells, and to activate CTLs in the glioma microenvironment (Figure 1E).

To test our therapeutic approach, we used the RCAS/t-va murine model to recapitulate low to high-grade glioma malignant progression in AYA mice (Suppl. Fig. 1A), as previously described [30, 31]. In brief, gliomagenesis is triggered by a right hemispheric parenchymal injection of DF1 cells for the release of PDGF RCAS viral particles in PTEN^KO^ pups (day 0-2 of age). This immunocompetent model supports gliomagenesis and tumor progression from low- to high-grade within 8 weeks (Suppl. Fig. 1A).

### GEMys-IL2 demonstrate stable expression of IL2

Three days post isolation and infection, mature myeloid cells were characterized with flow cytometry (Fig. 2A, Suppl. Fig. 2D). We evaluated the expression of CD45, CD11b, CD11c, MHC-II, CD115, Gr-1 (Ly6C/G), F4/80 in GEMys-EV or GEMys-IL2 using a gating strategy reported by Chow et al. [36]. We identified Dendritic Cells DCs (CD45^+^CD11b^+^CD11c^+^MHC-II^+^), Macrophages (CD45^+^CD11b^+^F4/80^+^Gr-1^-^), Monocytes (CD45^+^CD11b^+^Gr-1^lo^CD115^+^) and CD45^+^CD11b^+^Gr-1^hi^CD115^+^), Eosinophils (CD45^+^CD11b+Gr-1^-^CD115^int^F4/80^+^) and Neutrophils (CD45^+^CD11b^+^Gr-1^+^CD115^-^). The highly heterogeneous mature myeloid population in culture was mainly composed of DCs (1.46±0.47%), Macrophages (11.43±1.46%), Monocytes (24.40±6.33%), Neutrophils (37.11±3.61%), Eosinophils (less than 0.40%), and a 25.40% of uncharacterized cells. Among the uncharacterized cells, Mast cells and Megakaryocytes were clearly visible at the microscope in every batch of GEMys we generated (data not shown), but were not detectable by flow cytometry. Three days post isolation and infection, the aggregate GEMys-IL2 population showed a significant upregulation of IL2 gene expression (Fig. 2B) and IL2 secretion in the culture media (Fig. 2C). Unexpectedly, engineered IL-2 secreting myeloid cells consistently expressed and secreted more IFNγ than the GEMys-EV cells (Suppl. Fig. 2A,B), and this indirect effect might support the activation of tumor-infiltrating immune cells (Fig. 1A) [37]. To select the infected clones, cells were cultivated in selection media for 10-14 days (GEMys media enriched with puromycin 1μg/ml) prior to the engraftment in mice. To confirm the stable expression of IL2 and the clone selection, GEMys were monitored weekly by quantitative real-time PCR (RT-qPCR), flow cytometry, and fluorescent microscopy (Fig. 2D-F). Of note, the cells in selection media did not change their myeloid composition.

**Figure 2.**
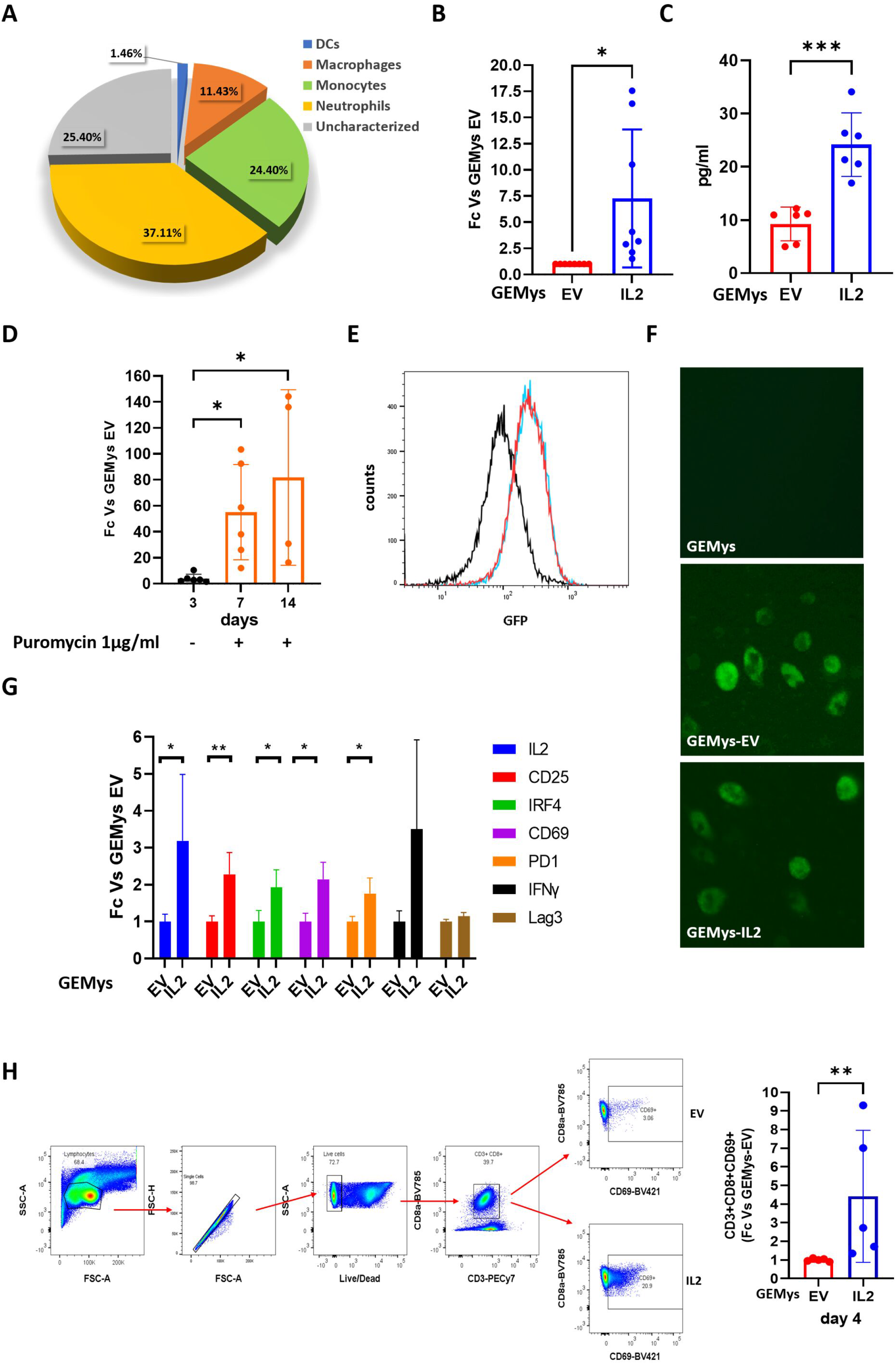
Generation of bone marrow-derived mature myeloid cells, engineered for the release of IL2 for the treatment of LGGs in vivo, clone selection and activation of primary murine CD8+ T cells by GEMys-IL2 in vitro. **A)** Flow cytometry evaluation of the GEMys myeloid composition in vitro (n=6). **B)** RT-qPCR analysis of the IL2 expression in vitro in GEMys expressing the empty vector (EV), or -IL2 at day 3 post lentiviral (LV) transduction. Results are expressed as fold-change of IL2 in GEMys-IL2, as compared with GEMys-EV (n=8). **C)** Quantification by ELISA of the IL2 released in the supernatant of GEMys in culture for 3days post LV infection (n=6). **D)** The stable expression of IL2 was reached after one week of clone selection in culture media supplemented with 1μg/ml of puromycin. RT-qPCR of IL2 gene expression in GEMys-EV, and GEMys-IL2. Results are expressed as the fold-change of IL2 in GEMys-IL2 cells in comparison with GEMys-EV (n=5). To confirm the positive selection of the transduced cells, the GFP positivity was evaluated by **E)** Flow cytometry (n=3), and **F)** Immunofluorescence analysis (n=3). Images of the green fluorescence have been acquired with 20X of magnification. **G)** CD8+ T cells were isolated from spleen of healthy syngeneic mice, and co-cultured with GEMys-EV, or -IL2 (T cells: GEMys ratio 4:1) for 24h. To analyze the T cells transcriptome, CD8+ cells were isolated by CD8+ magnetic negative selection before RNA extraction and RT-qPCR for the investigation genes involved in the T-cell activation (IRF4, PD-1, IL2, IFNγ,CD69, CD25a), or exhaustion (Lag3). **H)** FACS evaluation for CD69 expression in CD8+ T cells after 4 days of co-culture with GEMys-IL2, or with controls (GEMys-EV) (n=3). Unpaired two-tailed student’s t test calculated statistical significance. *P<0.05, **P<0.01, ***P<0.001.

### GEMys-IL2 activate primary murine CD8+ T cells in vitro

Cytotoxic T lymphocytes lead the adaptive immune anti-cancer response, and their activation is critical to effective immunotherapies. To test the ability of GEMys-IL2 to activate the target cells in vitro, we co-cultured primary murine CTLs with GEMys for 24h, and evaluated the canonical markers associated with cytotoxic T cell activation by quantitative real-time PCR (RT-qPCR) [38, 39]. We also evaluated the expression of Lag3, one of the primary markers associated with T cells inactivation/ exhaustion [40] in the same experiment. The co-culture of murine CTLs with GEMys-IL2 triggered the significant upregulation of IRF4 (2-fold), PD-1(1.8-fold), CD69 (2.1-fold), CD25a (IL2 receptor, 2.3-fold) and IL2 (3.2-fold), as compared with the co-culture of CTLs with control GEMys-EV (Fig. 2G). Notably, Interferon gamma (IFNγ) expression showed the same trend (3.5-fold increment), although not significant. No difference in Lag3 expression was detected in T cells co-cultured with GEMys-EV or IL2, thus confirming the absence of T cells exhaustion/inactivation in the experimental setting.

To further validate the activation of CD3+CD8+ cells in vitro, we performed 2 additional experiments. First, we co-cultured the CTLs with GEMys for 4 days, and evaluated the expression of CD69 by flow cytometry (Fig. 2H). CD69 protein expression measured in CD3+CD8+ T cells cocultured with GEMys-IL2 (ratio 10:1) was on average 3.3-fold higher compared to the levels in T cells treated with control GEMys-EV, hence confirming the results from CD69 gene expression (Fig. 2G). Second, we cultured GEMys for three days (80% of confluency) in the absence of puromycin, then harvested their media and used it to cultivate CTLs for 48h. Although the culture conditions were not ideal for the maintenance of primary T cells, the evaluation by RT-qPCR of T cell activation markers showed a significant upregulation of PD-1 (3.9-fold) and IRF4 (1.6-fold) in T cells cultured in GEMys-IL2 media, compared with T cells in GEMys control media (GEMys-EV, Suppl. Fig. 2C). Once again, the T cells inactivation marker Lag3 was not differentially expressed in CTLs cultivated in media isolated from GEMys-EV or -IL2, suggesting a supportive effect from secreted factors released by GEMys-IL2 on the activation status of CD3+CD8+ T cells.

### GEMys-IL2 are recruited and infiltrate the local glioma microenvironment

Using pre-clinical LGG to HGG murine models, we tested the efficacy of GEMys-IL2 as a novel cell-mediated immunotherapy. We engrafted 4-week-old LGG bearing animals with GEMys (Suppl. Figure 1). The RCAS glioma model also conferred the expression of luciferase in LGG cancer cells during tumor progression [31]. Therefore, we evaluated the tumor burden at day 25 by measuring the intracranial emission of photons released by the cancer cells using in vivo imaging system (IVIS) to randomize the mice into three treatment groups. Each group was engrafted at day 28 by retro-orbital intravenous injection of PBS (vehicle), 5×10^6^ of GEMys-EV, or 5×10^6^ of GEMys-IL2 cells. Mice were monitored post-treatment daily to evaluate toxicity or stress induced by the treatment. The systemic delivery of syngeneic myeloid cells was well tolerated by the animals, who showed no sign of stress, discomfort, or behavioral issues during the experiments. At 5 days post-engraftment, peripheral blood samples and brain tissues were isolated for further evaluation. We quantified the amount of IL2 in serum by ELISA, and verified that the amount of circulating IL2 in the animals engrafted with GEMys-IL2 was significantly higher than in animals treated with PBS, or with GEMys-EV (Fig. 3A).

**Figure 3.**
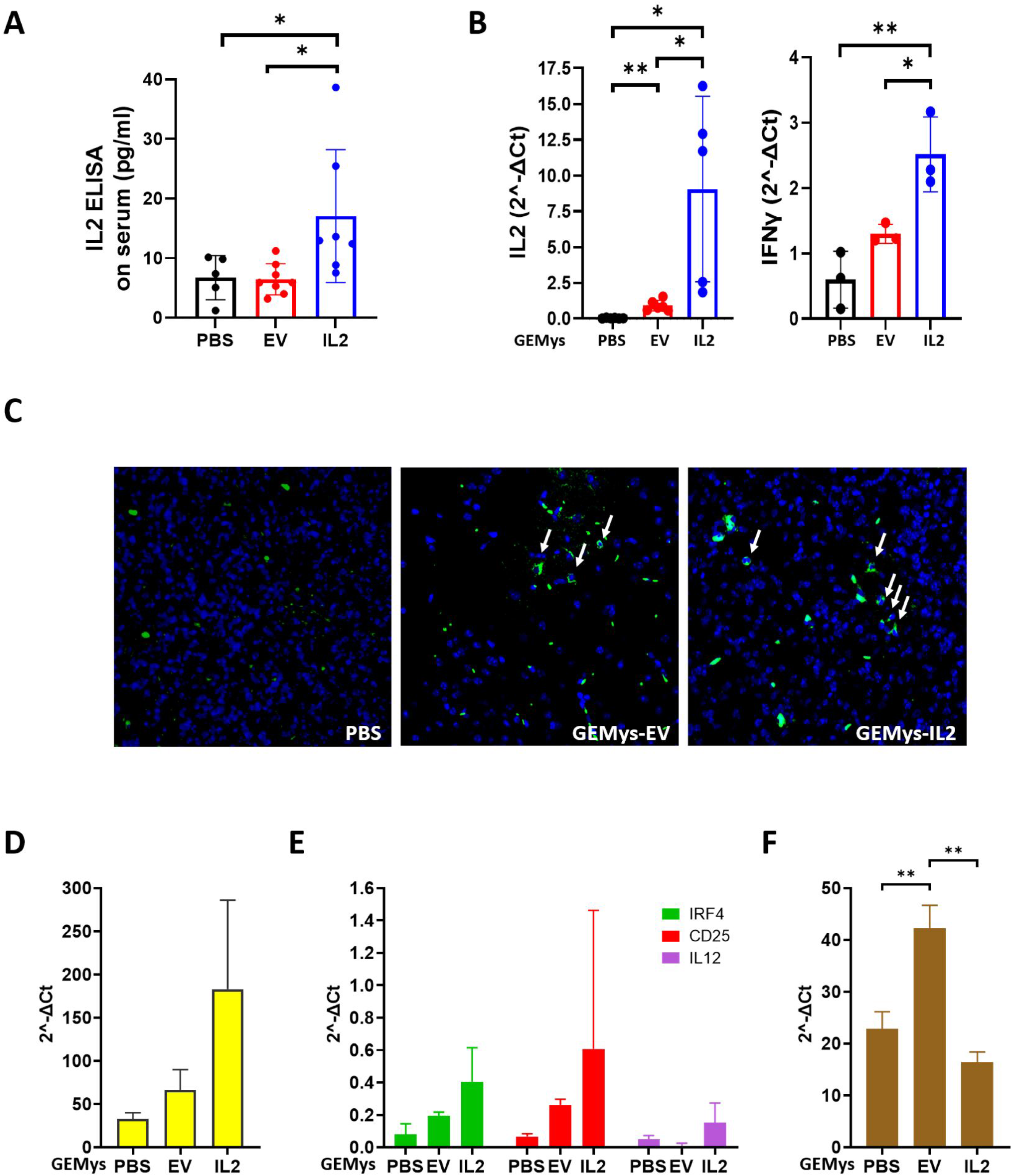
The treatment of RCAS/t-va LGG animals with GEMys-IL2 modify the tumor infiltrating immune cells transcriptome. LGG animals were treated for 7 days by retro-orbital intravenous (IV) injection (80μl) of GEMys-IL2, GEMys-EV, or vehicle (PBS). **A)** ELISA quantitation of circulating IL2 in the peripheral blood (n=6). **B)** RT-qPCR for the IL2 and IFNγ gene expression in immune cells infiltrating the TME. Brain tissues were (n=5). **C)** Immune fluorescence of brain sections of RCAS/t-va LGG animals, for the evaluation of the infiltration of GFP+ myeloid cells (**Green**). Brain sections were stained with Hoechst (**Blue**). (n=3). **D-F)** RT-qPCR for the evaluation of the expression of **D)** STAT1, **E)** IRF4, CD25, IL12 and **F)** LAG3 in infiltrating immune cells (n=3). Unpaired two-tailed student’s t test calculated statistical significance. *P<0.05, **P<0.01.

Previously, we demonstrated the recruitment and mobilization of peripheral bone marrow-derived myeloid cells to the glioma TME and therefore we hypothesized that systemically injected GEMys may home to the local glioma microenvironment [41]. To that end, we evaluated whether the GEMys circulating in the peripheral bloodstream of engrafted animals infiltrated the local the glioma TME after treatment using confocal fluorescent microscopy to demonstrate the presence of GFP-positive cells in brain coronal sections. Although this analysis required the application of algorithms to filter out the GFP cellular signal due to artifacts and basal levels of autofluorescence (distinctive of myeloid cells), we observed infiltration in the glioma microenvironment of GFP-positive myeloid cells in animals engrafted with GEMys-EV and GEMys-IL2 (Fig. 3C, Suppl. Fig. 2E). These results demonstrates that circulating GEMys pass the blood brain barrier (BBB) and localize to the TME. No GFP-positive cells were found in brain sections from control LGG animals injected with vehicle (Fig. 3C).

### GEMys-IL2 regulate immune cell transcriptomes in the glioma microenvironment

To assess whether the GEMys-IL2 influence the immune cell transcriptomes, we isolated the immune cells infiltrating the glioma microenvironment, and measured the gene expression of IL2 and IFNγ to evaluate the activation status of the T lymphocytes and myeloid cells [37, 42]. The analysis revealed that immune cells isolated from animals engrafted with GEMys-IL2 had a significant increase of IL2 (10-fold) and IFNγ (2.5-fold) expression compared with mice engrafted with GEMys-EV, or vehicle (Fig. 3B). To better characterize the activation status of the immune cells in the TME, we also investigated the expression of genes with a role in CD3+CD8+ T cell activation (STAT1, IRF4, CD25) [38, 39], or with T cell exhaustion (LAG3) [40](Fig. 3D-E). Here, RT-qPCR demonstrated the increase of STAT1, IRF4, CD25 (IL2 receptor), IL12, IL2, and IFNy expression (Fig. 3D-E) in immune cells of the TME isolated from animals engrafted with GEMys-IL2, in comparison to control GEMys-EV, or vehicle (PBS) TMEs. Remarkably, GEMys-IL2 also induced a significant reduction of Lag3 expression in vivo, suggesting an activated status of the effector T cells after treatment with GEMys (Fig. 3F, Table 2, Suppl. Fig. 3B). The heterogeneity of immune cell subpopulations and variable tumor burden in animals belonging to the same treatment groups contributed to the high standard deviation and intra-group variability of the experiment of Figure 3D-E, impacting statistical significance. Of note, the stimulation of immune cells mediated by IL2 release also induced the upregulation of STAT1 gene expression (Fig. 3D), which is crucial for the activation and recruitment of CD8+ T cells, and for the inhibition of infiltrating myeloid suppressive cells in solid tumors [43]. These results support our hypothesis that in vivo GEMys have a therapeutic effect, and in particular, that IL12 was upregulated in immune cells after treatment in vivo with GEMys-IL2 (Fig. 3E). In solid tumors, the upregulation of IL12 in the TME has been associated with T and NK activation, reprogramming of immunosuppressive myeloid cells, and improved survival [29, 44]. In summary, the treatment of LGG mice with GEMys-IL2 induced the upregulation of genes associated with M1-macrophage polarization, CTLs activation, and recruitment.

**Table 2.**
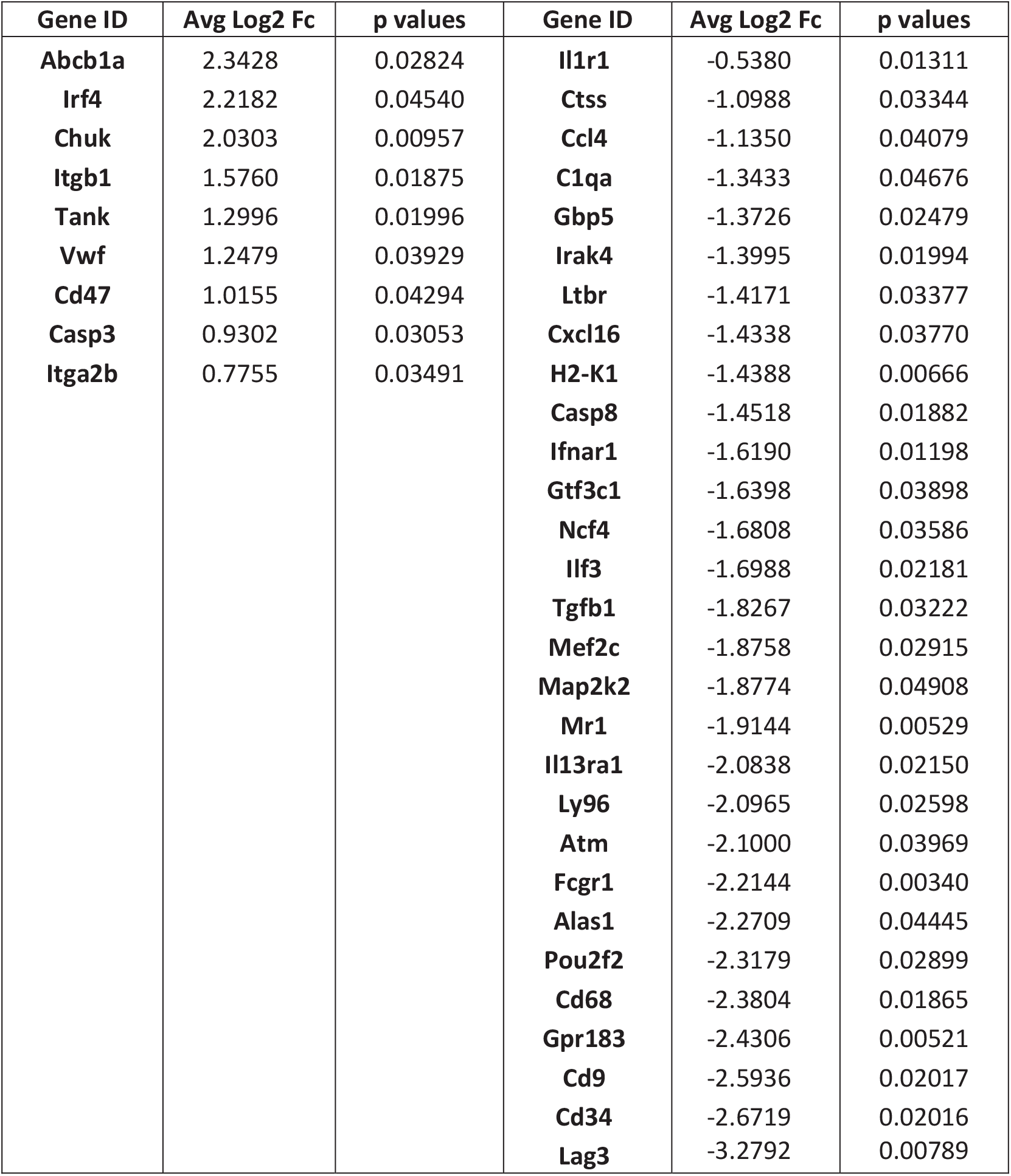
Comparison of the nCounter cancer immunology profile of immune cells infiltrating the glioma microenvironment, in vehicle or GEMys-IL2 engrafted animals. 5 days post engraftment of LGG mice with vehicle (PBS) or GEMys (IL2), and the immunological profile in the transcriptome of immune cells was analyses by nCounter. The gene list is organized by fold change (Log2 Fc). p-values were calculated by unpaired two-tailed T-test.

To corroborate our findings, we compared the RNA-based nCounter profile of gene expression from immune cells in the glioma microenvironment isolated from animals at day 5 post engraftment with GEMys-IL2, to immune cells from the TME of control animals injected with vehicle. Unsupervised hierarchical clustering identified that 38 genes that were differentially expressed among the two groups of animals analyzed (Table 1 and Suppl. Figure 3A,B). In immune cells infiltrating the TME of LGG animals engrafted with GEMys-IL2, the analysis of the top 10 upregulated genes identified that IRF4, which is associated with effector T cell activation, had significantly increased expression. Conversely, the top downregulated gene was Lag3. In addition, the Ingenuity Pathway Analysis (IPA®, Qiagen) predicted the upregulation of IL2, STAT5A, and STAT6 (Suppl. Figure 3C). Those genes are involved in the JAK/STAT pathway, associated with T cells activation and proliferation [45]. The software also identified the downregulation of FOXP3, the master regulator of the immunosuppressive activation of infiltrating Tregs [46]. The gene enrichment analysis of these significantly dysregulated pathways pointed out the upregulation of several nuclear molecules of the T cell receptor signaling pathway, involved in vital processes such as proliferation, activation, differentiation, CD8+ expansion, and polarization (Suppl. Fig 3D, p-value 0.00149). Moreover, the T cell exhaustion signaling was predicted to be one of the top downregulated pathways (Suppl. Fig. 3E, p-value 0.00004). Taken together, these findings demonstrate that the homing of GEMys engineered for the stable release of IL2 within the tumor microenvironment was associated with reprogramming of the immune cells toward an activated phenotype.

### GEMys-IL2 alter the immune cell composition of the glioma microenvironment

Our determination that GEMys-IL2 infiltrate the TME and efficiently secrete IL2 to reprogram the gene expression of immune cells in a murine LGG model, did not elucidate whether they were capable of potentiating the trafficking of cytotoxic T cells. To address this issue, we engrafted LGG animals with GEMys-IL2, and characterized the composition of the immune cells isolated from the glioma microenvironment 7 days post-engraftment by flow cytometry (Fig. 4A). As before, the negative controls were immune cells isolated from mice engrafted with GEMys-EV or with vehicle. RCAS/t-va animals treated with GEMys-IL2 or GEMys-EV showed an increased recruitment of CD45+ immune cells into the TME (Fig. 4B). In addition, the amount of CD45+GFP+ cells in mice treated with GEMys-IL2 was 2-fold higher than in animals treated with control GEMys (Fig. 4B).

**Figure 4.**
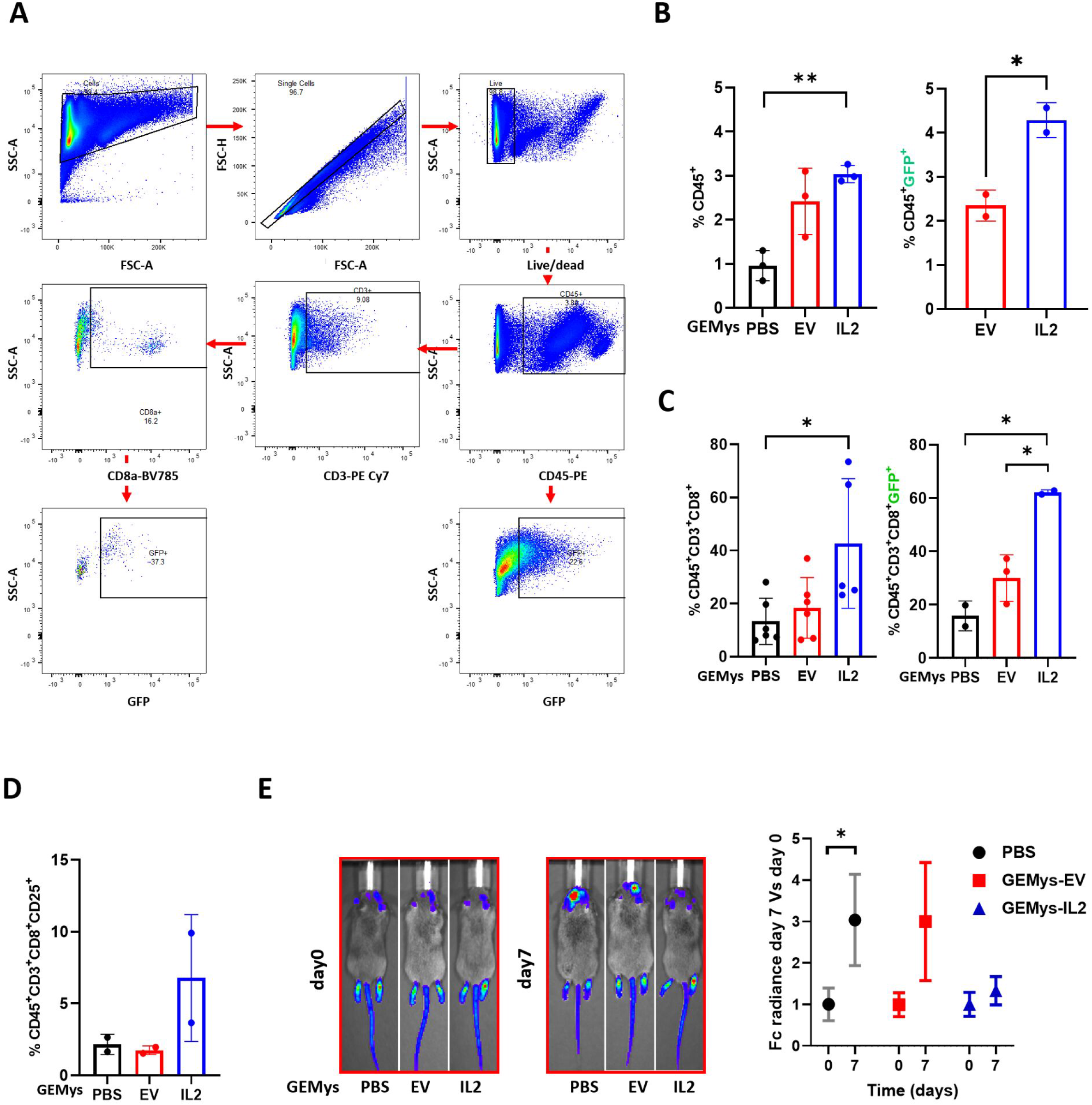
The treatment of RCAS/t-va LGG animals with GEMys-IL2 modify the composition, and activation of tumor infiltrating immune cells. LGG animals treated for 5 days by retro-orbital intravenous (IV) injection (80μl) with 5×10^6^ GEMys-IL2, GEMys-EV, or vehicle (PBS), and differences on the immune cell composition and activation in the TME were investigated by FACS. **A)** Flow cytometry gating strategy adopted for the analysis. Data were reported as % of investigated cell population as compared with the total amount of viable cells. **B)** % of CD45+ (left panel) and CD45+GFP+ (right panel) cells. **C)** % of CD45+CD3+CD8+ (left panel) and CD45+CD3+CD8+GFP+ (right panel) cells. **D)** % of CD45+CD3+CD8+CD25+ cells. **E)** Left panel: Tumor burden evaluation by IVIS of glioma progression in animals at day 0 and at 7 days post treatment. Right panel: quantification of the light emission (radiance) from cancer cells by IVIS (n=6 for PBS and GEMys-EV, 9 for GEMys-IL2). Data are expressed as fold-change of radiance compared to the average of radiance/treatment measured at day 0. Unpaired two-tailed student’s t test calculated statistical significance. *P<0.05.

One of the aims for this study was to demonstrate that pro-activating and bone marrow-derived myeloid cells engineered for the release of Interleukin-2 within the TME were able to recruit and activate anti-glioma cytotoxic T lymphocytes and arrest the tumor progression. In support of our hypothesis, FACS analysis of the immune cells revealed a significant 3-fold increased recruitment of cytotoxic T cells into the glioma microenvironment of animals treated with GEMys-IL2, as compared with controls (Fig. 4C). Of note, FACS demonstrated the upregulation of CD25, a known activation marker for cytotoxic T cells (Fig. 4D). Notably, cytotoxic T cells also expressed GFP, as explained by the fact that the GEMys-IL2 secreted IL2 molecules fused with GFP at the C-terminal end. Therefore, these data suggest the interaction of IL2-GFP with the IL2 receptor (CD25) on the target T cells is supported by the GFP positivity of CD3+CD8+ T lymphocytes.

### GEMys-IL2 homing to the glioma TME delays malignant progression

To investigate whether the treatment of LGGs in vivo had an impact on the tumor burden, we employed the RCAS/t-va murine model. We engrafted GEMys-IL2 in randomized LGG animals at week 4 post gliomagenesis as previously described, and we evaluated the tumor burden by IVIS post initial GEMys inoculation and at seven days post-treatment. As shown in Figure 4E, animals that received GEMys-IL2 had a remarkable reduction in tumor progression compared with control animals (GEMys-EV and PBS). To understand whether the reduced tumor burden was associated with an improved survival in vivo, we conducted a survival study and followed the tumor progression of these animals weekly by IVIS. As shown in the Kaplan-Meier survival curve, a single injection of five million syngeneic GEMys-IL2 cells was sufficient to induce a significantly prolonged survival in RCAS/t-va LGGs animals compared to animals injected with vehicle (PBS, p=0.0006), or to animals injected with GEMys-EV (p=0.0238) (Fig. 5A,B). Longitudinal IVIS monitoring demonstrated that, after an initial reduction of tumor progression in animals treated with GEMys-IL2 (Fig. 5B), tumor relapse began two weeks post-treatment. Indeed, circulating IL2 levels in serum at the endpoints were very similar in animals treated with a single dose of GEMys-IL2, GEMys-EV, or vehicles (Fig. 5C). Moreover, at the endpoints, all the animals had unequivocal transformation to HGG (Fig. 5D), therefore suggesting the effect of a single GEMys-IL2 treatment does not last longer than 7-14 days (Fig. 5B-D) although it delayed malignant progression significantly. In addition, no GFP-positive myeloid cells were found in the TME of animals treated with GEMys-EV or -IL2 at the endpoints (data not shown). Similar to the results of the in vitro (Fig. 1D-E, 2F) and in vivo (Fig. 3B,D-E, 4C) experiments, one dose of syngeneic treatment with GEMys-EV had a minimal therapeutic effect on the survival compared with animals that received vehicle. This was attributed to localization of naïve GEMys-EV (Fig. 3C, 4B) in the TME and their basal expression and release of growth factors and pro-activating cytokines for T cells, such as the Interleukin-2 and Interferon-Y (Fig. 3B).

**Figure 5.**
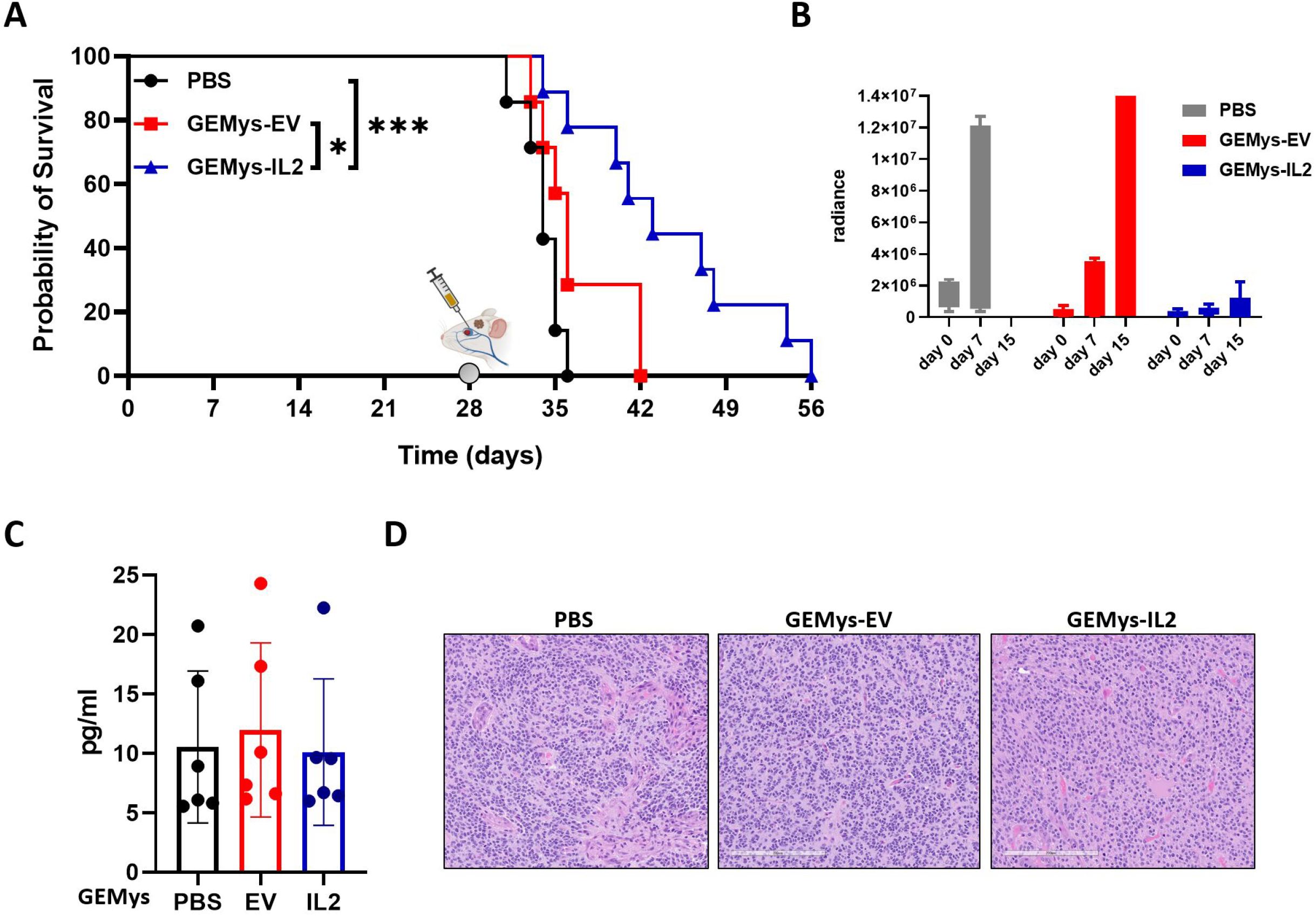
RCAS/t-va LGG animals treated with GEMys-IL2 improved the survival. **A)** Kaplan-Meyer evaluation of the survival of RCAS/t-va LGG animals, treated at day 28 by retro-orbital intravenous (IV) injection (80μl) of 5×10^6^ of GEMys-IL2, GEMys-EV, or vehicle (PBS). Log-rank Mantle-Cox test calculated the statistical significance. *P<0.05, *** P<0.001. **B)** Weekly evaluation of the tumor burden by IVIS, starting from the day of the treatment with GEMys, or vehicle. **C)** ELISA quantitation of circulating IL2 in the peripheral blood at the endpoints (n=6). **D)** H&E staining of 5μm paraffin embedded brain sections of animals at the endpoints (n=6).

## DISCUSSION

Here we report for the first time the successful use of a novel cell-mediated innate immunotherapy in a pre-clinical LGG to HGG progression model to delay glioma malignant progression. This novel immunotherapeutic approach consisted of the treatment of LGG glioma bearing animals with syngeneic bone marrow-derived myeloid cells, engineered for the stable release of Interleukin-2 (GEMys-IL2) into the glioma microenvironment. In the past, myeloid cells were considered just therapeutic targets in cancer, but new studies are revisiting their activity as potential therapeutic agents. To that end, two recent studies demonstrated the in vivo effect of primary derived and engineered human myeloid cells for solid tumors. Specifically, a study published in 2020 tested the efficacy of treating ovarian cancer in vivo with human macrophages engineered to bind of Her2 (anti-HER2 CAR-Ms) [47]. The second study from 2021 demonstrated the therapeutic efficacy of treating pre-metastatic lung cancer in a murine model with bone marrow-derived myeloid cells, engineered for the release of the pro-inflammatory cytokine IL12 [29].

The GEMys we generated in vitro were comprised of different types of mature myeloid cells and activated primary CD3+CD8+ T cells. Upon characterization, these myeloid cells were comprised of neutrophils, monocytes, and macrophages, but it remains to be determined whether the anticancer and pro-inflammatory activity in vitro and in vivo was mediated by a defined myeloid cell population or by diverse transduced myeloid cells. In addition, the treatment was delivered intravenously and well tolerated in an immunocompetent syngeneic LGG murine model. Recently, we proved that systemically injected bone marrow cells in glioma animals were able to pass the BBB and be recruited into the TME [41]. In a similar fashion, this study demonstrates that GEMys injected intravenously in immunocompetent glioma-bearing mice were recruited and infiltrated the local tumor microenvironment.

Once in the TME, the GEMys-released IL2 and our data suggests these cells and the secretion of IL2 influence the activation and composition of the infiltrating immune cells. Five days post-treatment with GEMys-IL2, we observed increased infiltration of immune cells in the LGG TME, in addition to increased trafficking and activation of cytotoxic T cells. This treatment effect was associated with pro-inflammatory reprogramming of the infiltrating immune cells in vivo as demonstrated by the upregulation of pro-inflammatory cytokines for T and myeloid cell activation, which aligned with a pro-inflammatory and activated status (IRF4, CD25, IL12, STAT1, IL2 and IFNγ) and the downregulation of the exhaustion marker Lag3. The immune cells recruited to the TME, and stimulated by the GEMys-IL2, upregulated STAT5A, STAT6 expression, which together with IL2 are all elements of the JAK/STAT pathway. This signaling cascade regulates T helper, CD3+CD8+ T, and NK cell activation and plays a crucial role in adaptive and innate immunity [45]. While historically the activation of the JAK/STAT pathway has been associated with tumor progression and reduction of anti-tumor immunity, recent investigations have suggested a more nuanced scenario, highly dependent on the specific cell type, especially in infiltrating myeloid cells [48].

In addition, evaluating the number of GEMys infiltrating the TME 5 days post-treatment with respect to the number of cells systemically injected is challenging due to cell state plasticity within the TME. However, one single dose of GEMys in LGG mice was sufficient to arrest glioma malignant progression and to significantly improve overall survival, but it was not curative. Upon survival study endpoint, we observed neither GFP positive myeloid cells in the TME of animals treated with GEMys-IL2 compared with control animals nor an increased amount of IL2 in the peripheral blood. Our results suggest that the GEMys effect does not last longer than 7 to 14 days in the treated animals. However, it remains unclear if the GEMys disappear by physiological turnover or are reprogrammed to be immunosuppressive after recruitment into the TME, or whether the prolonged stimulation of T cells with IL2 is causing exhaustion or inactivation. Future studies will address these fundamentally important questions. In addition, functional evaluation of the spatial distribution of other innate and adaptive immune cells will be critical.

In previous investigations using a similar glioma immunocompetent model, tumors were treated by injection of mature macrophages engineered for the release of IL12. However, the effect on the transcriptome was reached exclusively by intratumoral injection, and no survival data on the model was reported [49], hence the clinical relevance of the findings appear to be unclear. In fact, although intratumoral delivery of drugs may be feasible in more accessible solid tumors, the translation of such therapy to the clinic is still challenging for brain tumors [50].

This study is the first to demonstrate the pre-clinical activity of bone marrow-derived mature myeloid cells engineered to express IL2 to delay malignant progression of low-grade gliomas and prolong the OS. Due to a well-tolerated treatment course in our pre-clinical models with GEMys, and the positive effect on the recruitment of effector T cells, we believe the treatment may be more potent if the animals are treated with multiple administrations of GEMys or in combination with checkpoint inhibitors. These preliminary studies highlight a potential novel tool in developing a next-generation innate systemic immunotherapy, to be translated into the clinical arena.

## AUTHOR’S CONTRIBUTION

AC, MN, SR, CS, AH, HW, PF performed experiments. AC contributed substantially to develop the project, design the experiments, and to analyze data. GN contributed the nCounter data analysis and interpretation. AC, MN, SR, AH and DT interpreted the data. AC wrote the manuscript. ESM and TPC edited the manuscript. AC, MN, SR, CS, AH, GN, DT, HW, PF, ESM, TPC, PR were involved in the final editing and PR approved the final manuscript.

## ACKNOWLEDGMENT

We gratefully acknowledge Lara Rizzotto for carefully editing the manuscript, the OSU Genomics core facility (CCSG: P30CA016058) for the acquisition of the samples for the analysis of the transcriptome by nCounter (Nanostring), and Yang Hu for the scRNAseq data normalization, elaboration, and repository. We also greatly appreciate Ian’s Friends Foundation for grant support for our studies.

---

**Supplementary Table 1.**
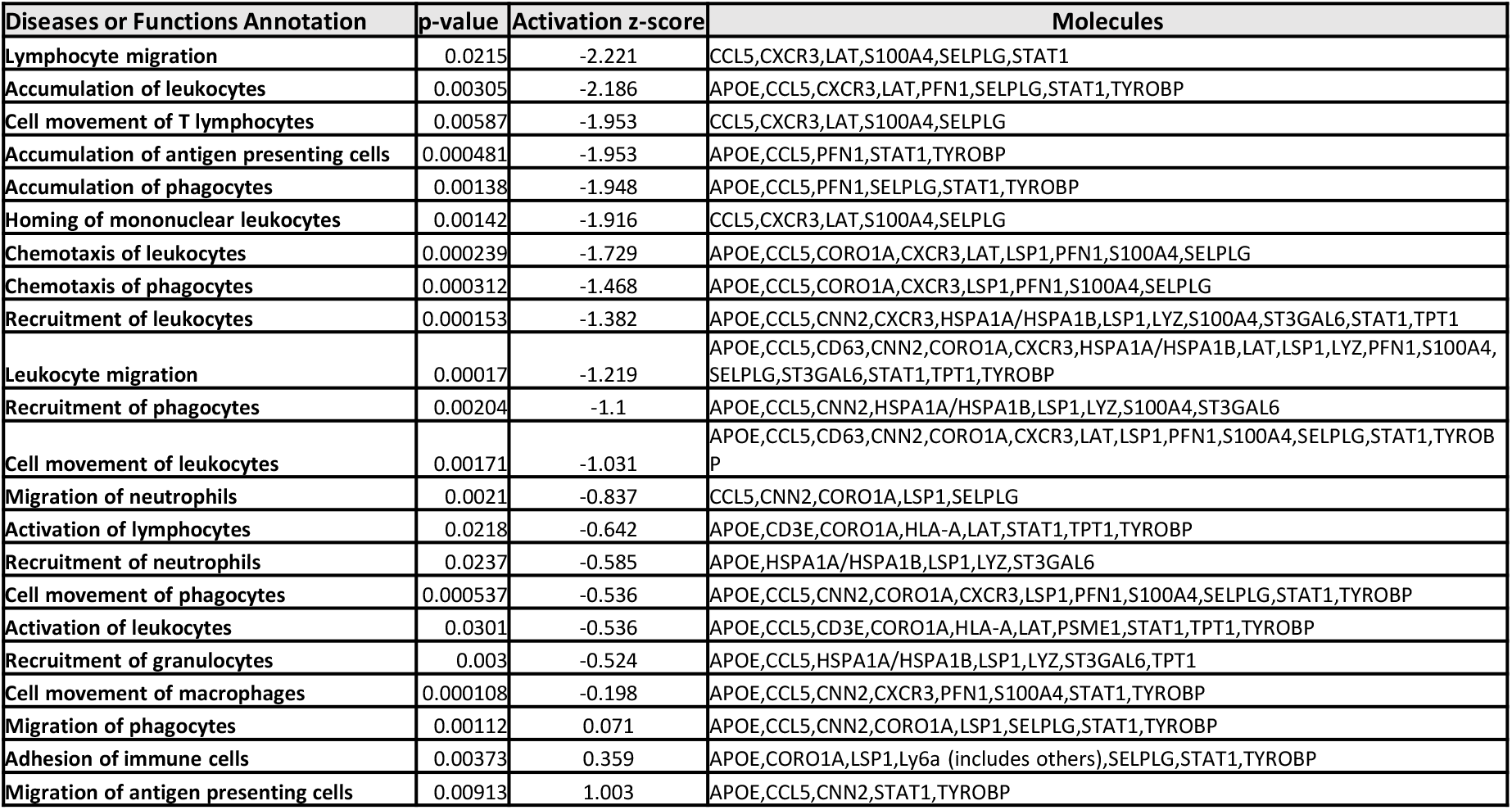
IPA Disease and biological function annotations of the deregulated genes involved in cell trafficking of TILs at the HGG, compared with TILs at the LGG.

**Supplementary Table 2.**
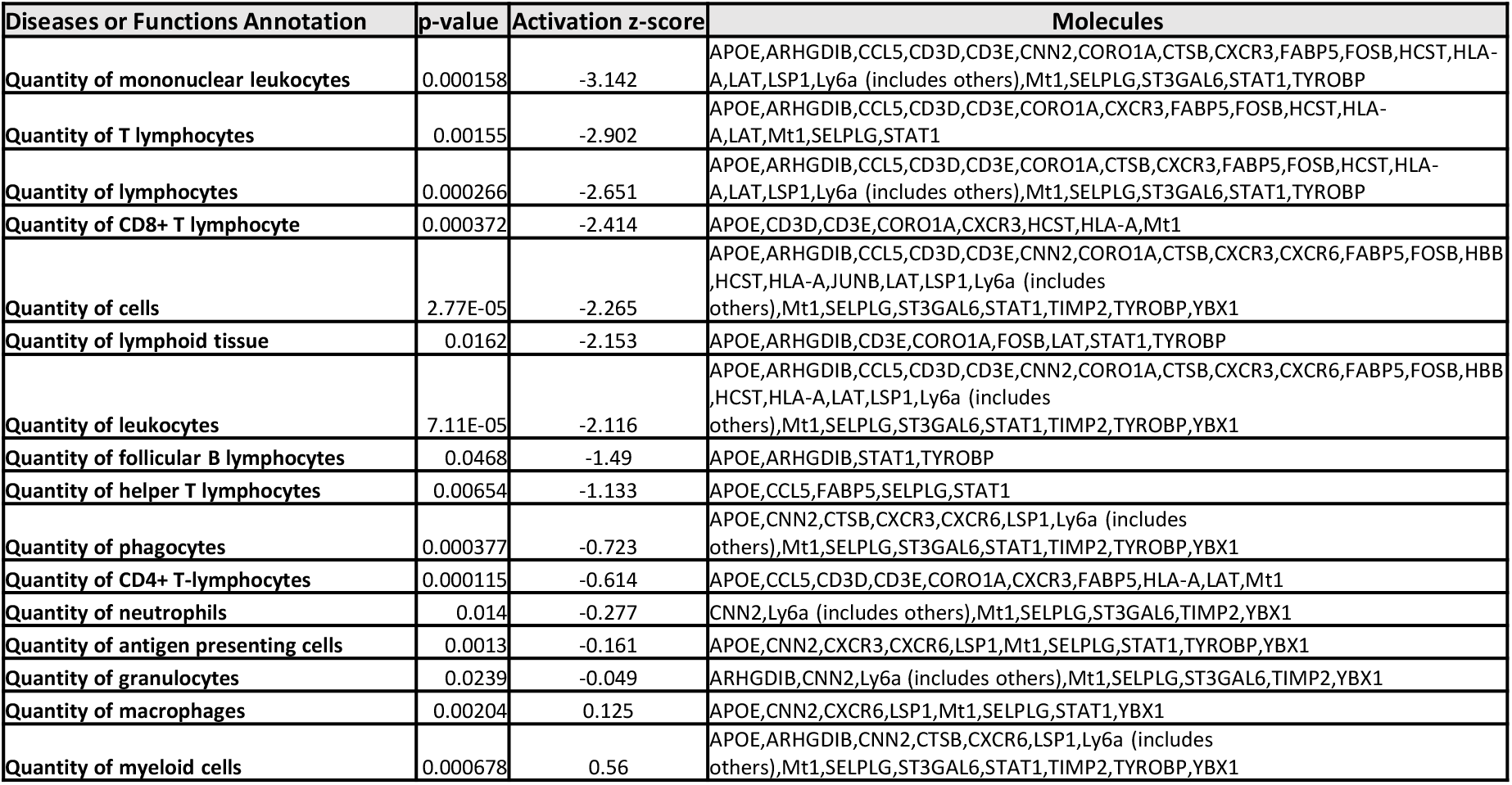
IPA Disease and biological function annotations of the deregulated genes involved in tissue morphology of TILs at the HGG, compared with TILs at the LGG.

**Supplementary Figure 1.**
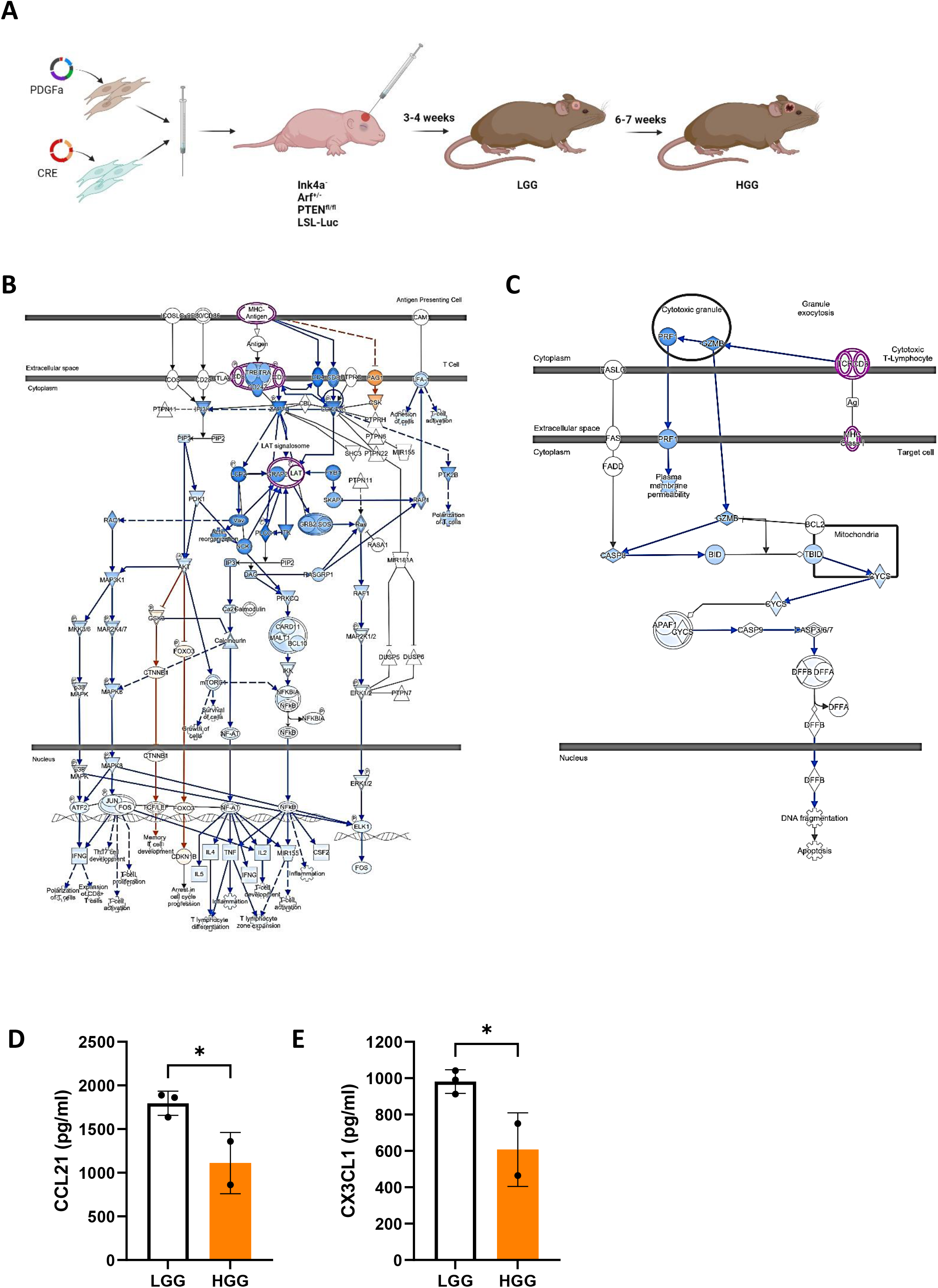
**A)** Graphic representation of the induction of gliomagenesis in RCAS/t-va model (Biorender.com). **B-C)** Evaluation by IPA of the regulation in TILs isolated from RCAS/t-va LGG, and HGG animals (n=3), and processed for scRNAseq, of the **B)** T cell receptor signaling and **C)** Apoptosis mediated by cytotoxic T cells signaling pathways. **D)** Analysis by cytokine array of the intratumoral protein expression of CCL21 in RCAS/t-va LGG, and HGG animals (n=2-3). **E)** Analysis by cytokine array of the intratumoral protein expression of CX3CL1 in RCAS/t-va LGG, and HGG animals (n=2-3). Unpaired two-tailed student’s t test calculated statistical significance. *P<0.05.

**Supplementary Figure 2.**
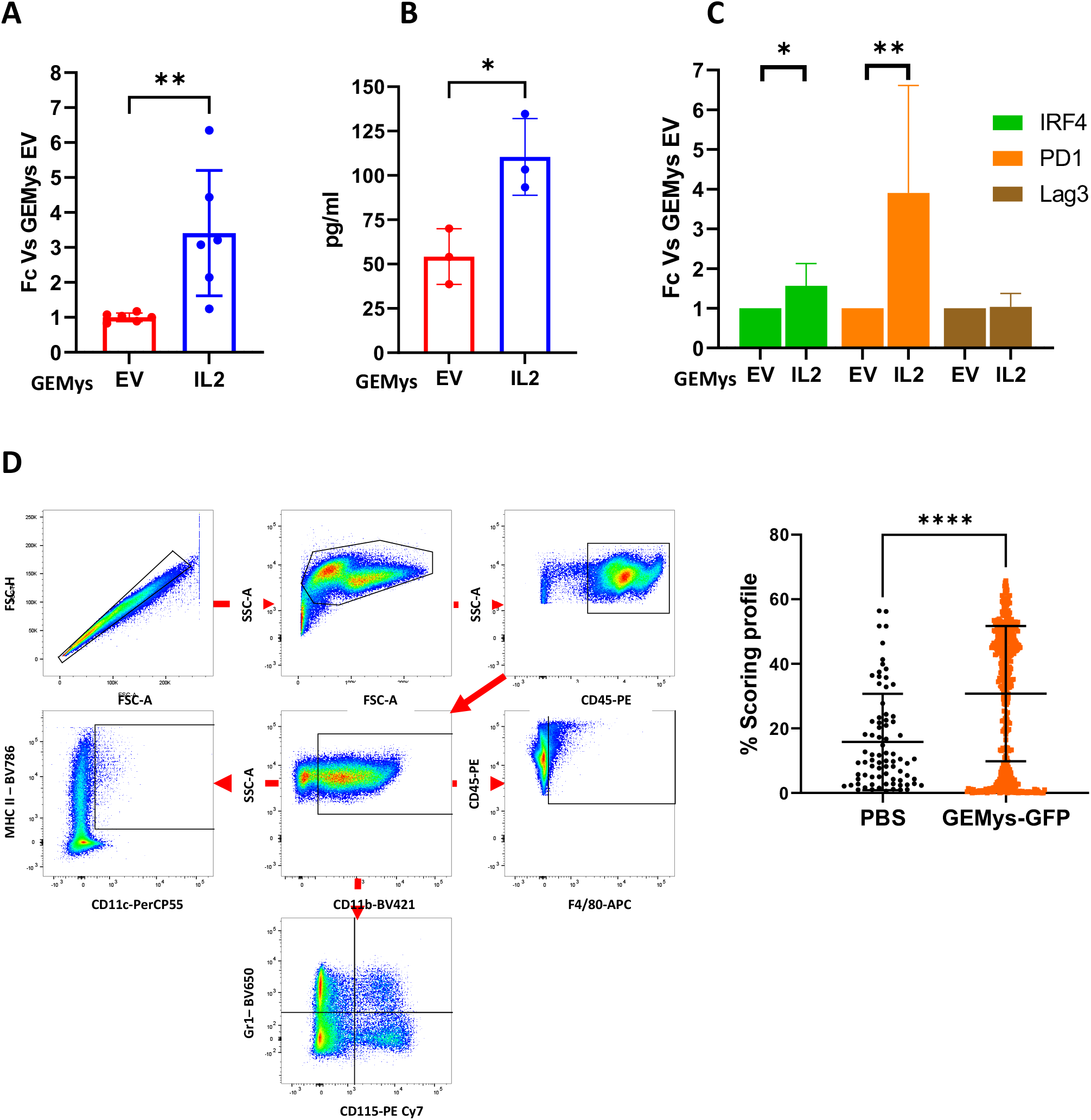
**A)** RT-qPCR of IFNγ gene expression in GEMys-EV, and GEMys-IL2. Results are expressed as the fold-change of IFNγ in GEMys-IL2 cells in comparison with GEMys-EV (n=6). **B)** Quantification by ELISA of the IFNγ released in the supernatant of GEMys in culture for 3days post LV infection (n=3). **C)** GEMys at 80% of confluency were cultivated in puromycin negative media for 3 days. GEMys-depleted and filtered media was used to cultivate for 48h CD8+ primary murine T cells, followed by RT-qPCR for the evaluation of the IRF4, PD-1 Lag3 gene expression. **D)** Flow cytometry gating strategy for the identification in GEMys in vitro of the myeloid cells composition. (n=6). **E)** IXMC-based molecular imaging analysis of GFP+ cells quantification in brain sections of LGG animals at day 5 post-engraftment with vehicle (PBS) or GEMys-GFP (n=3). Dots represent the GFP intensity colocalized with nuclear staining in 78 (PBS) and 443 (GEMys-GFP) microscopic fields. Unpaired two-tailed student’s t test calculated statistical significance. *P<0.05, **P<0.01, ****P<0.0001.

**Supplementary Figure 3.**
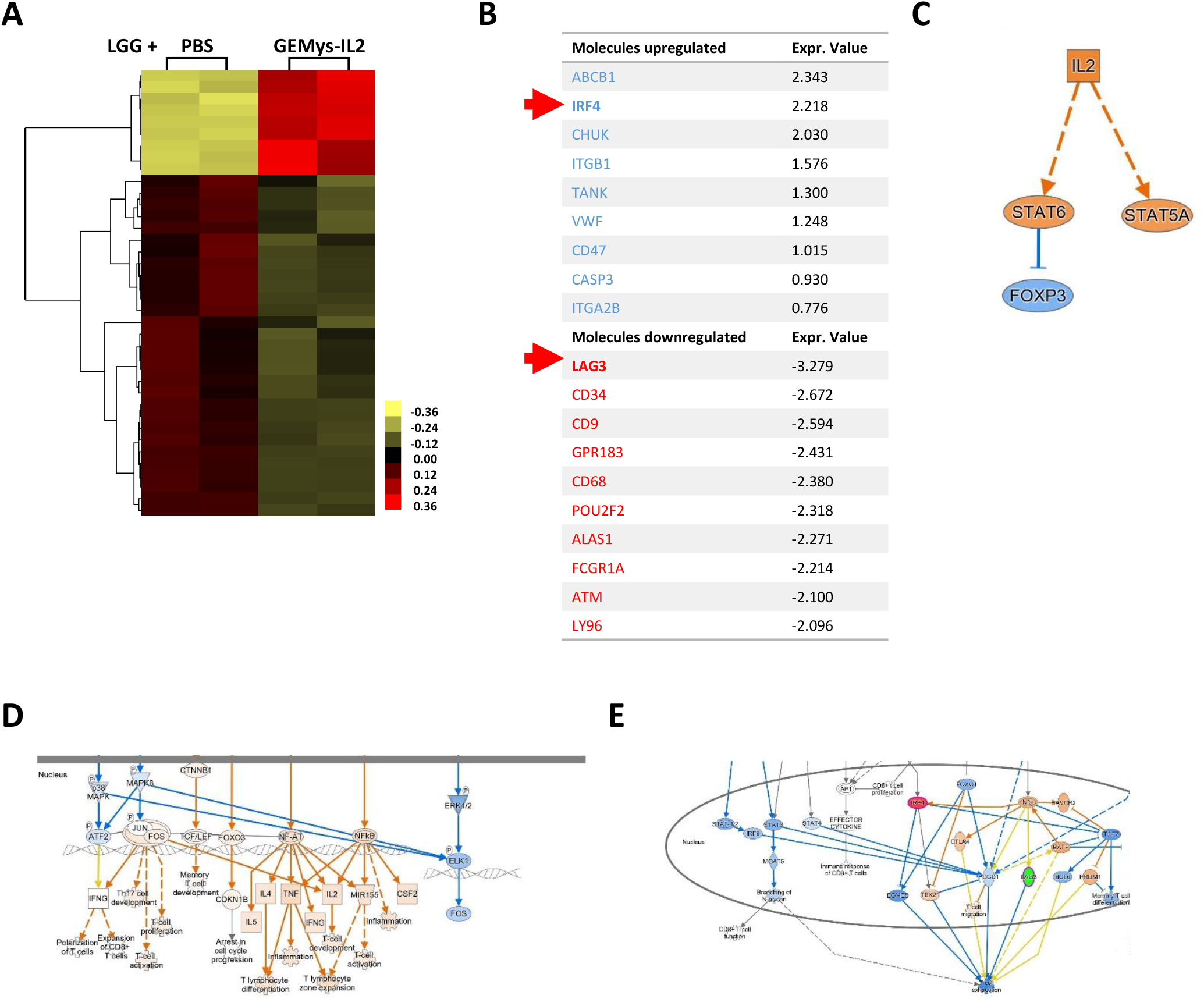
nCounter analysis of RCAS/t-va LGG animals treated for 5 days with GEMys-IL2, or vehicle (PBS). **A)** Dendrogram of the unsupervised hierarchical clustering analysis of nCounter®) mouse PanCancer Immune profiling assays in intratumoral immune cells (n=2). **B)** List of the significant top 10 upregulated (above) and of the top 10 downregulated (below) genes in the infiltrating immune cells of animals treated with GEMys-IL2, compared with animals treated with vehicle. Unpaired two-tailed student’s t test calculated statistical significance of genes with p<0.05. **C-E)** The signature was analyzed by IPA to estimate **C)** the predicted upstream targets, **D)** the gene expression regulation of the nuclear members of the T cell receptor signaling pathway, and **E)** the gene expression regulation of the nuclear members of the T cell exhaustion pathway.

