## SUPPLEMENTARY MATERIALS for "Genetically modified IL-2 bone marrow-derived myeloid cells reprogram the glioma immunosuppressive tumor microenvironment"

**Isolation of immune cells infiltrating the Glioma Microenvironment.** Animals were euthanized 5 days post treatment, brains were removed, and the area around the tumor was isolated from the rest of the right hemisphere. The brain tissue was mechanically dissociated on a 70µm cell strainer and collected in DPBS supplemented by the 10% of FBS (Corning). Cells were washed (DPBS+10% FBS), centrifuged at 500g for 10 minutes at +4°C, resuspended for 5 minutes on ice with RBC buffer (Biolegend) to lyse the red blood cells. RBC solution was quenched with 5 volumes of DPBS+10% FBS, and cells isolated by centrifugation. The stratification of the cells in 25% Percoll (GE Healthcare) was used to deplete the myelin layer and performed at 500g for 20 minutes at 18°C, in absence of break. Pellet was washed in DPBS+10% FBS and stained for flow cytometry or processed for RNA isolation and analysis of the transcriptome.

**Single cell RNA sequencing (scRNAseq).** Tumor or normal tissue was excised from the brain's right hemisphere, at the injection site. The tissue was then dissected into smaller pieces, digested with 2 mg of Papain (Brainbits) for 20 minutes at 37°C, and filtered through a 70 µm cell strainer. CD45+ cells were isolated by magnetic positive selection (Miltenyi). The database generated for the IPA analysis is publicly available (weillcornellmed.shinyapps.io/3_samples_ShinyCell/)

**CD8+Tcells co-culture with GEMys.** Spleens were harvested from 4-week-old NTV-a;Ink4a$^{-}$Arf$^{+/-}$;PTEN$^{fl/fl}$;LSL-Luc mice without tumors, and immediately mechanically dissociated on a 70µm cell strainer (Thermo Scientific). Cells were washed in PBS and processed for CD8 positive selection (StemCell Technologies). For investigation by flow cytometry, cells were co-cultured in 24 well plates with GEMys-EV, or GEMys-IL2 with a T-cells : GEMys ratio of 1:4 for 4 days in RPMI media supplemented with 10% FBS (Gibco). Cells were harvested, stained, and acquired as described above. To profile the transcriptome, CD8+ cells were co-cultured with GEMys-EV, or GEMys-IL2 with a ratio of 4:1 for 24hs, followed by CD8+ T cells magnetic negative selection (StemCell Technologies), total RNA extraction and real-time PCR. Flow cytometry was used to evaluate the protein expression, and for extracellular staining and acquisition, a protocol outlined by Patel et al was adapted for murine samples [1]. To evaluate the activation status of the cytotoxic T lymphocytes (CTL) from the GEMys co-culture, cells were stained with a panel of titrated antibodies including ZOMBIE NIR (Biolegend 423105 ), CD45-PE (Biolegend 103106), CD3-PECy7 (Biolegend 100319), CD8-BV786 (Biolegend 100749), CD25-BV605 (Biolegend 102035), and CD69-BV421 (Biolegend 104527). CTL were gated as viable CD45+ CD3+ CD8+ and their activation status was assessed by CD25 and CD69 with florescence minus one (FMO) control for each activation marker. As positive controls for activation, CTL were stimulated for 48hr with anti-CD3/28 Dynabeads (Thermofischer 11456D) as described by Patel et al. Stained cells were acquired on a LSR Fortessa (BD Biosciences), and analyzed using Flowjo software.

1. Patel, T., et al., *Development of an 8-color antibody panel for functional phenotyping of human CD8+ cytotoxic T cells from peripheral blood mononuclear cells.* Cytotechnology, 2018. **70**(1): p. 1-11.

**Flow cytometry**

**GEMys Composition:** To determine the composition of GEMys in vitro, a titrated flow cytometry panel consisting of ZOMBIE NIR (Biolegend 423105), CD45-PE (Biolegend 103106), CD11b-BV421 (Biolegend 101235), F4/80-APC (Biolegend 123116), Gr1-BV650 (Biolegend 108441), CD115-PeCy7 (Biolegend 135523), CD11c-PerCP/Cy5.5 (Biolegend 117327), and IA/IE-BV786 (Biolegend 107645) was designed with FMO controls.

**Immune profile of the Glioma Microenvironment:** Immune cells were isolated as already described and stained with two titrated flow cytometry panels. One panel included the T cell activation markers previously described with the addition of CD335-APC (Biolegend 137608) for natural killer cells and another myeloid panel including ZOMBIE NIR (Biolegend 423105), CD45-PE (Biolegend 103106), CD11b-BV421 (Biolegend 101235), F4/80-APC (Biolegend 123116), and Gr1-BV650 (Biolegend 108441). Since both the EV and IL-2 GEMys express GFP, it was measured in the FITC channel.

**Quantitative Real Time PCR (RT-qPCR).** Total RNA from primary isolated murine cytotoxic T cells and myeloid cells was extracted using RNeasy mini kit (Qiagen), and total RNA from primary immune cells isolated in the glioma microenvironment was extracted using RNeasy micro kit (Qiagen). RNA quantity and quality were assessed by Nanodrop (Thermo Fisher), and cDNA was synthetized using High-Capacity cDNA Reverse Transcription Kit (Applied Biosystems) and in accordance with the manufacturer's protocol. SYBR Green RT-qPCR experiments were executed following the manufacturer's protocols (Applied Biosystems) using PrimeTime qPCR primer assays listed in Table 1 (IDT), and in a StepOnePlus system (Applied Biosystems).

| | Primer sequence | |
|---|---|---|
| **Gene ID** | **Forward** | **Reverse** |
| **GAPDH** | AATGGTGAAGGTCGGTGTG | GTGGAGTCATACTGGAACATGTAG |
| **TBP** | TGTATCTACCGTGAATCTTGGC | CCAGAACTGAAAATCAACGCAG |
| **SDHA** | TCCATACACCGAATAAGAGCAAA | ACCAGCCCTAGTGACCAT |
| **CD69** | ACGGAAAATAGCTCTTCACATCT | ACCACTATTAACACAGCCCAAG |
| **FOXP3** | CTGGTCTCTGCAGGTTTAGTG | CTGTCTTCCAAGTCTCGTCTG |
| **IL2** | GCAGGATGGAGAATTACAGGAA | GCAGAGGTCCAAGTTCATCTTC |
| **CD25 (IL2ra)** | CTGCCTCTTCCTGCTCATC | GCTCTGACTTTTCTAGCTTGCT |
| **IL12a** | ACAGATGACATGGTGAAGACG | CTCTCGTTCTTGTGTAGTTCCA |
| **IFNγ** | CTGAGACAATGAACGCTACACA | TCCACATCTATGCCACTTGAG |
| **IRF4** | CTCTTTGACACACAGCAGTTTC | TCACCAAAGCACAGAGTCAC |
| **LAG3** | TCAATGCCACTGTCACGTT | GTTACCTCACACAACAGCTTC |
| **PD-1** | GTACCCTGGTCATTCACTTGG | ATTTGCTCCCTCTGACACTG |
| **STAT1** | TTGACAAAGACCACGCCTT | GACTTCAGACACAGAAATCAACTC |

**Supplementary Table 3.** mouse-specific real-time PCR primers used in the project.

**mRNA profiling.** Total RNA was extracted from the immune cells infiltrating the glioma microenvironment using RNeasy micro kit (Qiagen), and after 5 days post engraftment of LGG RCAS/t-va mice with vehicle (PBS), or GEMys-IL2 (n=2). RNA concentration and purity were assessed by Nanodrop (Thermo Fisher) and TapeStation (Agilent). The mRNA profiling of 770 genes involved in cancer immunology was evaluated by the nCounter® (NanoString) mouse PanCancer Immune profiling panel (NS_Mm_CancerImm_C3400). Gene profiling normalization was performed using nSolver™ Analysis Software (NanoString Technologies, Seattle, WA, USA), as recommended by NanoString. Negative controls were used to perform background subtraction. After technical normalization, the data were biologically normalized by calculating the geometric mean of the top 100 genes in all samples. P-values were calculated using the t-Student test. Genes with a p-values<0.05 were retained for the downstream analyses. Clustering analysis was performed with Cluster 3.0, and the heatmap was generated with Java Treeview. Ingenuity Pathway Analysis (IPA) was used to predict disease-related functions, upstream regulators, networks, and pathways affected by the tumor progression or by the treatment in vivo.

**Immunofluorescence.** Whole brains were dissected from mice at endpoint and fixed in 10% buffered formalin phosphate for 24 hours. 5µm paraffin-embedded brain tissue sections were obtained from mice treated with vehicle, GEMys-EV or GEMys-IL2 and euthanized 5 days post treatment. After deparaffinization and hydration, antigen retrieval was performed in citrate pH 6.0 buffer on a high-pressure setting. The sections were blocked with TBST with 10% normal goat serum for one hour at room temperature. Sections were treated with nuclear stain Hoescht (1:5000, Life Technologies) for 0 minutes at RT and mounted using Prolong gold anti-fade mounting media. 20X images was acquired with the ImageXpress Micro Confocal High-Content Imaging System (IXMC, Molecular Devices). To measure the infiltration of GFP positive cells in the TME, the percentage of positive cells to GFP and Hoescht colocalization were calculated by the multiwavelength scoring module of the MetaXpress software (Molecular Devices).

**Cytokine array.** Immune cells infiltrating the glioma microenvironment in the RCAS murine model were collected from the tumoral mass in the right hemisphere of brains from animals at week 3 (low-grade glioma), or at week 7 (high-grade glioma). Tumor progression was validated by histological evaluation of 5µm paraffin-embedded brain tissue sections stained by H&E. Total protein lysates were generated with RIPA buffer supplemented with HALT proteases inhibitor cocktail (Thermo Scientific), the protein concentration measured by BCA (Thermo Scientific), following the protocols of the manufacturer. The cytokine profile was investigated by Mouse Cytokine 44-plex Discovery Assay (Discovery Assay).

**Enzyme-Linked immunosorbent Assay (ELISA).** To perform the quantification of the circulating murine IL2 in the peripheral blood, or of the released IL2 in the culture media, we performed ELISA as described by the manufacturer (R&D Systems). All incubations were conducted at room temperature unless otherwise noted. Briefly, the plate was coated overnight with 100µL of primary antibody and blocked in 1% BSA for 1h. 100µL samples or standards were seeded in triplicate and incubated for 2h. The plate was washed, and biotinylated anti-IL2 was added to each well and incubated for 2h. 100µL Streptavidin-HRP was added and allowed to stand for 20 min. In the dark, 100µL of substrate solution was added to each well and incubated for 15 min. 50µL of stop solution was added and absorbance at 450nm and 570nm (wavelength correction) was read on a Synergy 2 microplate reader (Biotek).
